# Dihydroartemisinin inhibits EMT of glioma via gene BASP1 in extrachromosomal DNA

**DOI:** 10.1101/2023.03.13.532418

**Authors:** Que Zhongyou, Zhou Zhiwei, Liu Sheng, Zheng Wenhua, Lei Bingxi

## Abstract

The mechanism of dihydroartemisinin (DHA) inhibiting the migration and invasion of glioma in an ROS-DSB-dependent manner has been revealed. Extrachromosomal DNAs (ecDNAs) which are generated by DNA damage have great potential in glioma treatment. However, the role of ecDNAs in DHA’s pharmacological mechanisms in glioma is still unknown. In this study, DHA was found to inhibit proliferative activity, increase ROS levels and promote apoptosis in U87 and U251 cells. Migration and invasion have also been suppressed. ecDNA expression profiles were found in gliomas. EcDNA-BASP1 was found, by means of bioinformatics analysis, to be present in GBM tissues and positively correlated with patient prognosis. Proliferation, migration and invasion were upregulated after knockdown of ecDNA-BASP1. The expression of vimentin and N-cadherin also had the same tendency. Finally, we found that the ecDNA-BASP1 content in nude mouse transplant tumors was significantly increased after DHA treatment, which might exert a better suppressive effect on glioma. The upregulation of tumor suppressor ecDNA-BASP1 played an important role in the suppression of glioma progression induced by DHA. EcDNA-BASP1 may inhibit glioma migration and invasion through repressing epithelial–mesenchymal transition (EMT).

## 1. Introduction

Glioblastoma (GBM) is the most common malignant tumor of the central nervous system, with a median survival of only 8 months [1]. Nowadays, the main strategy of GBM treatment is a combination of surgery, radiotherapy, and chemotherapy [2]. Even though targeted therapy, immunotherapy, and electric field therapy have emerged in recent years, they still cannot fundamentally transform the prognosis of patients with GBM [3–5]. Epithelial–mesenchymal transition (EMT) is the process of cellular change from an epithelial phenotype to a mesenchymal phenotype. In glioma, EMT plays an important role, not only as a central link in the control of GBM migration and invasion, but also as a strong correlate of GBM cell stemness and other characteristics [6]. Verhaak et al. identified four clinically relevant subtypes of GBM by means of genomic analysis [7]. Recently, Cyril Neftel et al. revealed the plasticity of different subtypes of GBMs by means of single-cell-sequencing, which is similar to EMT [8]. Hadi Zarkoob et al. investigated the relationship between GBM molecular subtypes and EMT by bioinformatics. They found that molecular subtypes have similarity with the EMT process, and the mesenchymal subtype has the closest ties [9]. Therefore, exploring the molecular mechanisms regulating EMT is of great value in finding out potential molecular targets for the treatment of GBM.

Extrachromosomal DNA (ecDNA) is free circular DNA (extrachromosomal circular DNA, eccDNA) which is found in the cytoplasm outside of the nuclear genomic DNA and ranges from a few hundred to several thousand base pairs in size [10]. Although ecDNA was first discovered in the 1960s, the function of ecDNA has received attention only in recent years because of the improvement of bioinformatics [11,12]. Compared to chromosomal DNA, ecDNA is able to carry oncogenes more efficiently to participate in processes such as drug resistance, and even act as mobile enhancers to regulate tumor-associated gene transcription [13,14]. EcDNA was found in more than half of all tumors, and glioma is one of the tumors with the highest ecDNA content so far [15]. It is suggested that ecDNA may also play an important role in the EMT process of GBM.

Dihydroartemisinin (DHA) is a potent antimalarial drug, and the antitumor potential of DHA has also been recognized [16]. Current studies have found that the pharmacological mechanism of artemisinins is mainly related to reactive oxygen species (ROS) [17]. Our group found that DHA was able to inhibit the migratory and invasion of U87 and U251 cells in a ROS-dependent manner. In glioma cells, ROS production can eventually inhibit the expression and activity of MMP7/MMP9 in U87 and U251 by inducing double strand break (DSB) activation of p53 and the downstream β-catenin signaling pathway [18]. Since the mechanism of ecDNA generation is currently believed to be associated with DNA damage, we hypothesize that ecDNA plays an important role in the inhibition of glioma EMT by DHA [19].

In this study, we first verified that DHA could effectively inhibit malignant biological behaviors such as the proliferation, migration and invasion of U87 and U251. Subsequently, the changes to the ecDNA expression profile after DHA treatment were analyzed using second-generation sequencing. The bioinformatics and TCGA database analysis finally identified ecDNA-BASP1 as a key factor affecting EMT in U87 and U251 cells. BASP1 (ENSG00000176788) is the parental gene of ecDNA-BASP1, which is located in chr5:17228272-17228562 and was named by our group, and the function of ecDNA-BASP1 is not clear. This study reveals that ecDNA-BASP1 may play an important role in the EMT of glioma and provides a new target and strategy for future GBM drug therapy.

## 2. Materials and Methods

### 2.1 Cell culture

The U87 MG (RRID: CVCL_3429) and U251 MG (RRID: CVCL_A5HR) cell lines (human glioma cell lines) were obtained from Nanjing KGI Biotechnology Co. All of the glioma cells were cultured in Dulbecco’s modified Eagle medium (DMEM) containing 10% fetal bovine serum (FBS) (Hyclone, USA). The cell lines were cultured in a 37°C incubator with a humidified atmosphere containing 5% CO_2_.

### 2.2 Cell proliferation assay

Cell viability was measured using the CCK8 assay kit (Sigma, USA). We seeded cells into 96-well plates (Corning) at a density of 5×10^3^ cells with 200μL of medium per well and further cultured for 24 h. Then, each well was treated with 20 μL of CCK8 for another 1.5 h at 37 °C. Finally, an ELISA plate reader (SYNERGY4, USA) was used to record the optical density at 570 nm (OD570). Five independent experiments were carried out.

### 2.3 ROS detection

We made use of the ROS assay kit (Beyotime, Shanghai, China) to detect intracellular ROS according to the manufacturer’s instructions. Briefly, we seeded 5000 cells per well in a 96-well plate and cultured for 24 hours. Then, each well were incubated with 2’,7’-dichlorofluorescein-diacetate (DCFH-DA) for 1 hour. Next, each well was washed with serum-free medium three times. The level of ROS was imaged and analyzed with a fluorescence microscope (Olympus, Japan).

### 2.4 Cell apoptosis analysis by flow cytometry

The effect of DHA on apoptosis was assayed using an Annexin V-Fluorescein Isothiocyanate (FITC)/Propidium Iodide (PI) kit (Beyotime Ltd., Shanghai, China). First, cells were exposed to 50μM DHA for 24 h and collected. Then, the cells were resuspended with the use of binding buffer and added to 5 μL of Annexin V-FITC and then 5 μL of PI for a 15-min incubation in darkness. Analyses of the results were performed by means of flow cytometry (BD Biosciences).

### 2.5 Mitochondrial membrane potential (MMP) detection

MMP was detected by the fluorescent probe JC-1 (mlbio Co.). After treated with DHA (50 μM) for 48h, cells were cultured with JC-1 (10 μM) at 37°C for 30 min. Then they were washed by PBS. Finally, the fluorescence intensity of JC-1 was detected by means of fluorescent microscopy (Olympus, Japan).

### 2.6 Cell migration ability detected by wound scratch assay

Groups of cells were inoculated onto six-well plates and three sets of replicate wells were set up. After cell fusion reached approximately 90%, six-well culture plates were drawn with 20 μL perpendicular to the pre-prepared horizontal lines. The plates were washed with PBS and incubated in 1% FBS DMEM medium three consecutive times. The Ibidi culture system was also used in our experiments [20]. This was repeated three times for imaging measurements. Data were analyzed by Image Pro Plus 6.0.

### 2.7 Transwell migration and invasion assay

The migration ability of U87 and U251 cells was assessed through a transwell assay. Transwell plates (Corning Costar) placed in 24-well plates with a well size of 8 μm were used to culture U87 and U251 cells. Then, 500 μL of conditioned medium (supplemented with 10% of FBS) was added into the lower chamber as an migration-inducing agent. DHA at a concentration of 50 μM was added to the upper chamber. After 24 hours of incubation, we took out the transwell inserts and carefully removed the residual cells on the upper surface using cotton swabs. The migrating cells on the lower surface were washed gently with PBS, fixed with methanol and glacial acetic acid (3:1 mixture) for 30min, and then stained with Giemsa stain for 15 min.

When it came to transwell invasion assays, Matrigel (BD Biosciences, USA) was added to the upper surface of the transwell chambers and diluted with culture medium at a ratio of 1:7. Other steps were the same as the transwell migration assay. Finally, the average number of migrated cells was counted using Image Pro Plus 6.0 software.

### 2.8 eccDNA enrichment for circle-Seq

High-throughput sequencing of Circle-Seq eccDNA was provided by Shanghai Cloud Sequence Biotechnology Co. Cell precipitates were resuspended in L1 buffer (Plasmid Mini AX; A&A Biotechnology) supplemented with proteinase K (ThermoFisher) and digested overnight at 50 °C. The digested samples were subjected to alkaline treatment and column purification according to the instructions of the Plasmid Mini AX kit. The column-purified DNA samples were digested by FastDigest MssI (ThermoFisher) for 16 h at 37 °C to remove the mitochondrial circular DNA. The treated samples were used as templates for the amplification of eccDNA by phi29 polymerase (REPLI-g Midi Kit). The amplified DNA was fragmented using a Bioruptor ultrasonicator and the fragmented DNA was indexed using the NEBNext^®^ Ultra II DNA Library Prep Kit. Libraries were sequenced in high throughput on a NovaSeq 6000 sequencer using 150bp double-end mode.

### 2.9 ecDNA Data analysis

Raw data were obtained from sequencing with an Illumina NovaSeq 6000 sequencer. Raw data quality control was first performed using Q30 values, using cutadapt software (v1.9.1) to de-join and remove low-quality reads to obtain high-quality clean reads. Clean reads were compared to the human reference genome (hg19) using bwa software (v0.7.12). Then, we identified eccDNA in all samples using circle-map software (v1.1.4), and the number of soft-clip reads overlapping with the break point was calculated as the raw count using samtools (v1.9) software, and edgeR (v0.6.9) software was used for standardization and the calculation of ploidy variation and the p-value between two groups/samples to screen differentially expressed eccDNA. Gene annotation of identified eccDNA and differential eccDNA was performed using bed tools software (v2.27.1). The annotated differential genes were used for GO function analysis and KEGG pathway analysis. eccDNA visualization was performed using IGV software (v2.4.10). All the analysis of ecDNA data in this research was provided by Shanghai Cloud Sequence Biotechnology Co [21].

### 2.10 Validation of eccDNA recordings

Real-time PCR was conducted using a QuantStudio 5 Real-Time PCR System (Thermo Fisher, US) and qPCR SYBR Green master mix(Cloud Sequence Biotechnology Co., Shanghai, China). PCR oligos were designed by Primer 5.0, and we prepared the Realtime PCR reaction system (2 × Master Mix 5 μL, 10uM PCR specific primer F 0.5μL, 10uM PCR specific primer R 0.5μL, added water to a total volume of 8μL). We mixed the solution by flicking the bottom of the tube and centrifuged briefly at 5000 rpm. We then added 8uL of the mixture to each well of the 384-PCR plate, and added a corresponding 2μL of the DNA template (100ng genomic DNA or 2μl of digested DNA). We carefully adhered the sealing film sealing membrane and mixed by means of brief centrifugation. We then placed the prepared PCR plate on ice prior to setting up the PCR program, and placed the above PCR plate on a Realtime PCR instrument for PCR reactions. All indications were performed according to the following procedure: 95°C, 10 min; 40 PCR cycles. To establish the melting curve of PCR products, the amplification reaction was performed according to the following procedure: 95°C, 10 s; 60°C, 60 s; 95°C, 15 s, and then heated slowly from 60°C to 99°C. The target DNA and internal reference of each sample were subjected to real-time PCR reactions separately. The data were analyzed by the 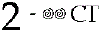 method.

### 2.11 Bioinformatic analysis

Gliomas RNA Seq data and clinical data were downloaded in the Cancer Genome Atlas Program (TCGA, https://portal.gdc.cancer.gov/), including 150 normal brain samples and 701 glioma samples. The differential analysis was carried out with statistical settings such as logFC filter = 1 and FDR filter = 0.05. The EMT-related markers were downloaded from the GESA database (https://www.gesa-ingenieure.de/). The volcano diagram, Venn diagram, univariate COX regression analysis, multivariate COX regression analysis, ROC curve analysis, heatmap analysis, and survival analysis were then conducted with R software. The above bioinformatic analysis involved Per software and R software, including different R packages such as Limma, Pheatmap, Survival, Survminer, SurvivalROC and Caret.

### 2.12 BASP1 knockout

The following sgRNA-primers were designed from http://www.rgenome.net/cas-designer/:

sgMOB1#1-oligo1 CACCGCTTAGGGAGGCAGACGTCAA
sgMOB1#1-oligo2 AAACTTGACGTCTGCCTCCCTAAGC
sgMOB1#2-oligo1 CACCGCATATTTGAATTAAGCAAAG
sgMOB1#2-oligo2 AAACCTTTGCTTAATTCAAATATGC
sgMOB1#3-oligo1 CACCGACACATTTACAGGTCATAGG
sgMOB1#3-oligo2 AAACCCTATGACCTGTAAATGTGTC

We then set up the digestion reaction (1.5 μg of pX459, 2 μL of 10X NEBuffer 2.1, 1 μL of BbsI (NEB), and added H_2_O to a final volume of 20 μL). We incubated the digestion reaction at 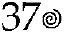 for at least 1h. 200 ng agarose gel was used to ensure complete digestion. We then added 1μL of CIP to the digestion reactions, incubating for 1h at 37°C. The gel purified the digested plasmid, and phosphorylated and annealed each pair of oligos. We then annealed the mixture in a thermocycler and set up the ligation reaction, and And hatched at room temperature for 10 minutes as recommended by the manufacturer. We then transferred 2-4μL of the final product into 50μL of competent cells. The selected colonies and sequences were verified with U6 sequencing (primer U6 seq F: ACTATCATATGCTTACCGTAAC).

### 2.13 Western Blot

The expression of vimentin and E/N-cadherin was detected by Western blotting. U87 and U251 cells were firstly washed with PBS three times and then lysed with ice-cold radio immunoprecipitation assay buffer which was supplemented with protease inhibitors. The cells containing lysate were incubated on ice for 30 minutes. Next, we used an ultrasonic crusher to disrupt the pellet, centrifuged the samples at 14,000g for 30 min at 4 °C, and finally collected and transferred supernatant to a new tube. We put the sample in a 99°C water-bath heater for 5 min to denature the proteins. Used BCA protein analysis kit (Beyotime Biotech Co. Ltd, China), and then subjected to electro phoresis with equal amounts (20–25 g) for each lane. The protein was separated in 10% SDS-polyacrylamide gel electrophoresis (PAGE), transferred into a PVDF membrane, and then incubated with rabbit polyclonal antibodies against vimentin (diluted at 1:1000) (Abcam Cat# 4211-1, RRID: AB_765097), N-cadherin (diluted at 1:1000) (BD Biosciences Cat# 610921, RRID:AB_398236), and E-cadherin (diluted at 1:1000) (SAB, 40860). A mouse polyclonal antiGAPDH antibody (diluted at 1:2000) (Hangzhou HuaAn Biotechnology Cat# EM1101, RRID:AB_2811078) was used as an internal control. After overnight incubation, the protein in PVDF membrane was washed with TBST, and then incubated with corresponding secondary antibodies. The protein bands were scanned with ChemiImager 5500 V2.03 software, and the protein expression was estimated by calculating the integrated density values with a computerized image analysis system (Fluor Chen 2.0)

### 2.14 Orthotopic implantation model in nude mice

All animal experiments were carried out in the lab of Gunagzhou Forevergen Co. and approved by the Animal Experiment Ethics Committee of Sun Yat-Sen University (LACUC number: L102022016000D). We bought the athymic BALB/C nude mice (4 weeks old, male) from the Cancer Institute of the Chinese Academy of Medical Science. We fixed each mouse with a brain stereoscopic locator after anesthetizing to prevent the head from shaking. We used a stereotaxic device to locate the caudate nucleus of the right brain, which was 3 mm from the cranial midline and 1 mm from the coronal suture. We drilled a hole and inserted the needle 6 mm into the skull, then withdrew 1 mm, and slowly injected 10 μL of U87 and U251 cells at a density of 1 × 10^5^ cells. We kept the needle inside for 5 min to prevent cells from overflowing. Mouse were then treated with a dose of DHA of 10 mg/kg, once a day for 45 days. The mice were killed on the 45th day after injection [22].

### 2.15 HE staining

All HE staining experiments were carried out by Shanghai Gefan Biotechnology Co. The brain tissues of mice were preserved in 10% formalin. The 10% formalin-fixed brain tissues were washed. Dehydrated in ethanol and soaked in xylene and melted paraffin and the tissues were cut into thin slices and stained with hematoxylin and eosin for the tumor areas.

Stained sections are photographed with a camera under an optical microscope (Olympus, Japan).

### 2.16 Immunohistochemistry assay

Immunohistochemistry(IHC) assay were performed to detect the expression of ki67 and Caspase 3. Washed the slides sequentially with specific reagents. Incubated in PBS (pH=7.4) containing 3% H2O2 for 5 to 10 minutes at room temperature to block endogenous peroxidase activity. Sections were rinsed with PBS for 5 minutes. Rinsed with PBS for 5 minutes after antigen retrieval. Applied blocking antibody and incubated for 30 minutes at room temperature which followed by shaking off residual liquid. Sections wre rinsed after incubation overnight at 4°C using an antibody of Ki67/Caspase 3 (dilution of 1:200). Sections were incubated at 37°C for 40 minutes using horseradish peroxidase-conjugated secondary antibody. Sections were rinsed and then stained with DAB stain. Rinsed and then stain the sections with hematoxylin.Finally,the stained sections were observed and photographed under a light microscope (Olympus, Japan).

### 2.17 In situ hybridization

Tissue chips and tissue sections from mice were dewaxed with xylene, rehydrated through an ethanol gradient and then treated with 3% H_2_O_2_ for 10 min. Subsequently, we treated the tissue samples with pepsin diluted in 3% fresh citrate buffer at 37°C for half an hour. After washing with PBS, pre-hybridization was performed using 20 mL of the pre-hybridization solution at 37°C for 2 h. Then, the hybridization was performed with DIG-labelled probes for overnight incubation at 37°C. Afterwards, the sections were treated in blocking solution for 30 min at 37°C after high stringency washes, and then incubated with alkaline phosphatase-conjugated sheep anti-DIG Fab fragments for 60 min at room temperature. Positive staining of ecDNA-BASP1 was observed by adding BM purple AP substrate according to the manufacturer’s instructions (Shanghai Gefan Biotechnology Co)

### 2.18 Clinical materials

Patient information and clinical outcomes were retrieved from the Sun Yat-sen Memorial Hospital. Before the operation, all patients provided written informed consent for their glioma samples to be used for research. Our tissue chips included a total of 60 patients, with 120 samples and paired data, including: 33 grade 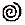 glioma samples and 33 normal control samples from the same patients, 15 grade 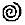 glioma samples and their controls, 11 grade 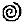 samples and their controls, and 1 grade 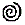 glioma samples and their controls. All of the clinical data were de-identified prior to analysis.

### 2.19 Statistical analysis

All data in accordance with normal distribution were described as mean ± S.D. All related statistical analysis was carried out with SPSS 20 software. We made use of Student’s t-test to assess the differences between two groups and the one-way analysis of variance (ANOVA) test to compare multiple groups followed by Dunnett’s post hoc test. P < 0.05 was considered to indicate statistical significance.

## 3. Results

### 3.1 Effects of DHA on proliferation, ROS, apoptosis and migration and invasion of U87 and U251 cells

First, we treated U87 and U251 cells with different concentrations of DHA (0μM, 50μM, 100 μM) for 24h and assayed them using the CCK-8 assay. We found that the proliferation of U87 and U251 cells was significantly inhibited at 50μM for 24h, while higher concentrations (e.g., 100μM) and time did not significantly increase the inhibition rate of glioma cells (Figure1 A). Considering that the effective concentration and time of action of DHA are closely related to the drug purity and the status of experimental cell lines, 50μM and 24h were used as the concentration and time point in the subsequent vitro experiments.

**Figure 1.**
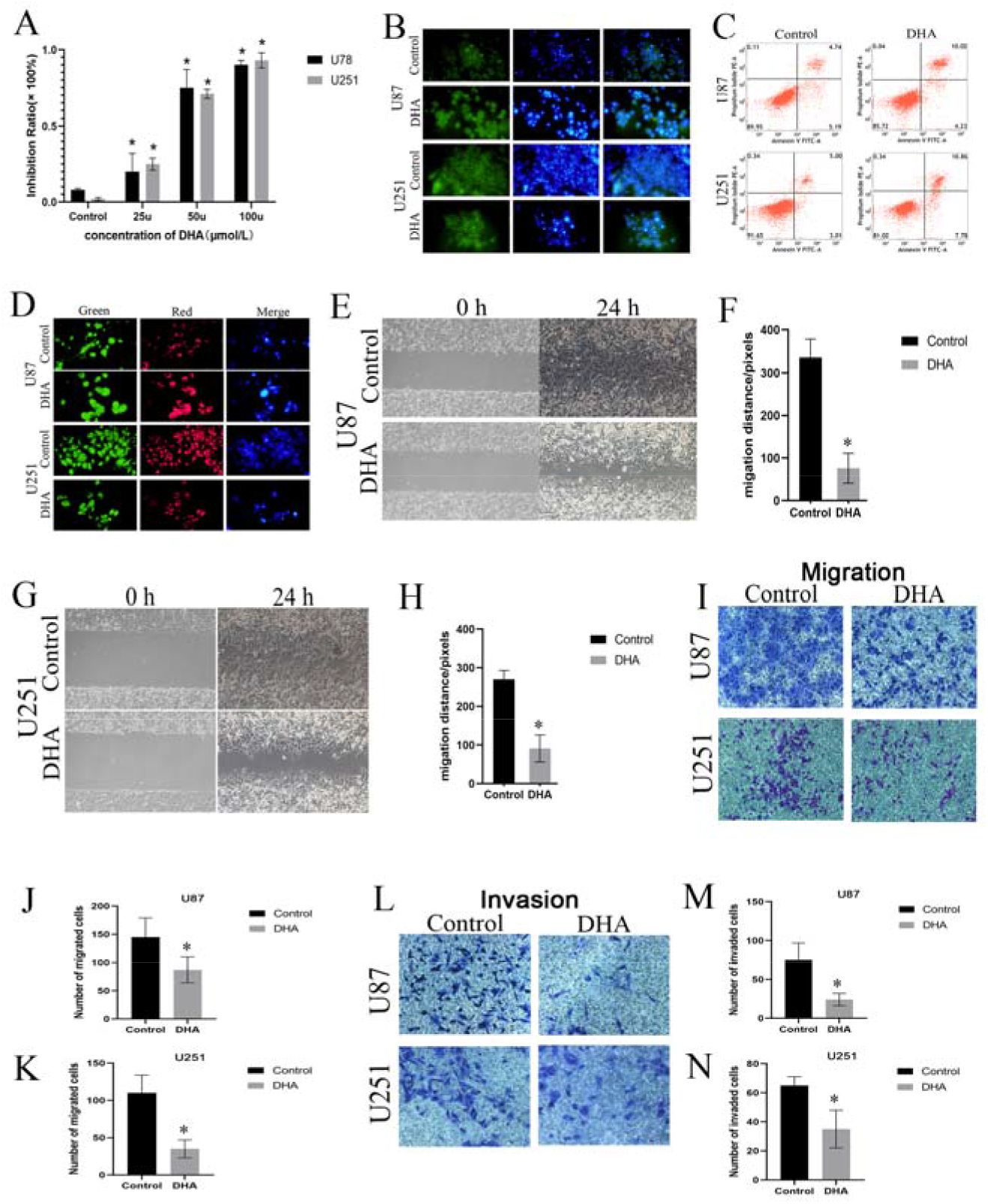
DHA inhibited proliferation, migration, and invasion and promoted ROS generation and apoptosis in U87 and U251 cells. **(A)** The inhibition ratio of U87 and U251 cells treated with DHA at different concentration (25μM, 50μM, 100μM) for 24h. **(B)** The change in ROS generation in U87 and U251 cells treated with DHA for 48h. **(C)** The apoptosis was activated by 50μM DHA treatment for 48h in U87 and U251 cells measured by flow cytometry. **(D)** The apoptosis was activated by 50μM DHA treatment for 48h in U87 and U251 cells measured by confocal microscopy. **(E)** Wound scratch assay showing the change in migration in U87 cells after treating with 50μM DHA for 24h. **(F)** Statistical results of the wound scratch assay in U87 cells. **(G)** Wound scratch assay showing the change in migration in U251 cells after treating with 50μM DHA for 24h. **(H)** Statistical results for the wound scratch assay in U251 cells. **(I)** Transwell migration assay in U87 and U251 cells after treating with 50μM DHA for 24h. **(J)** Statistical results for the transwell migration assay in U87 cells. **(K)** Statistical results for the transwell migration assay in U251 cells. **(L)** Transwell invasion assay in U87 and U251 cells after treating with 50μM DHA for 24h. **(M)** Statistical results for the transwell invasion assay in U87 cells. **(N)** Statistical results for the transwell invasion assay in U251 cells. *P<0.05

The changes in ROS in glioma cells after DHA treatment were subsequently detected by fluorescent probe. ROS was found to be significantly elevated in U87 and U251 cells with 50μM DHA for 24h (Figure1 B). As apoptosis is a common mechanism of many anti-glioma drugs, DHA has been found to exert anti-tumor effects by inducing apoptosis in glioma cells. We studied the effect of DHA on the apoptosis of U87 and U251 cells by using flow cytometry. Flow cytometry revealed that the apoptosis level of U87 and U251 cells was significantly increased after 50μM DHA action on U87 and U251 cells for 48h (Figure 1C), and the same fact was observed by fluorescence confocal microscopy (Figure 1D). Migration and invasion ability is an important cause of glioma progression, and it has been shown that DHA can significantly inhibit the migration and invasion ability of U87 and U251 cells. In the present study, we further verified the effect of 50μM DHA on the migratory invasion ability of U87 and U251 cells using an ibidi Culture-Insert 2 Well system and a transwell assy. We found that 50μM DHA could significantly inhibit the migration ability of U87 and U251 cells after 24h (Figure1 E-H). The transwell assay showed that the migration and invasion of U87 and U251 cells were significantly inhibited by 50μM DHA after 24h (Figure1 I-N).

### 3.2 Changes in ecDNA expression profile in U87 and U251 cells after DHA treatment

To investigate the changes to the ecDNA expression profile in U87 and U251 cells after DHA treatment, Circle-Seq technology was used. A large number of ecDNA fragments were detected in all four groups, including 731,745 ecDNAs in the U87-Control group, 113,255 ecDNAs in the U87-DHA group, 32,657 ecDNAs in the U251-Control group and 277,101 ecDNAs in the U251-DHA group. They were evenly distributed on each chromosome, and the size was mainly distributed between 0.01 and 10 kb (Figure2 A). Compared with the control group, a portion of ecDNA expression profiles were significantly changed in both U87 and U251 cell lines (Figure2 B). Based on the parental genes of the 1108 significant ecDNAs, GO functional analysis and KEGG pathway enrichment analysis were performed to further investigate the biological functions of the changing ecDNA expression profiles.

**Figure 2.**
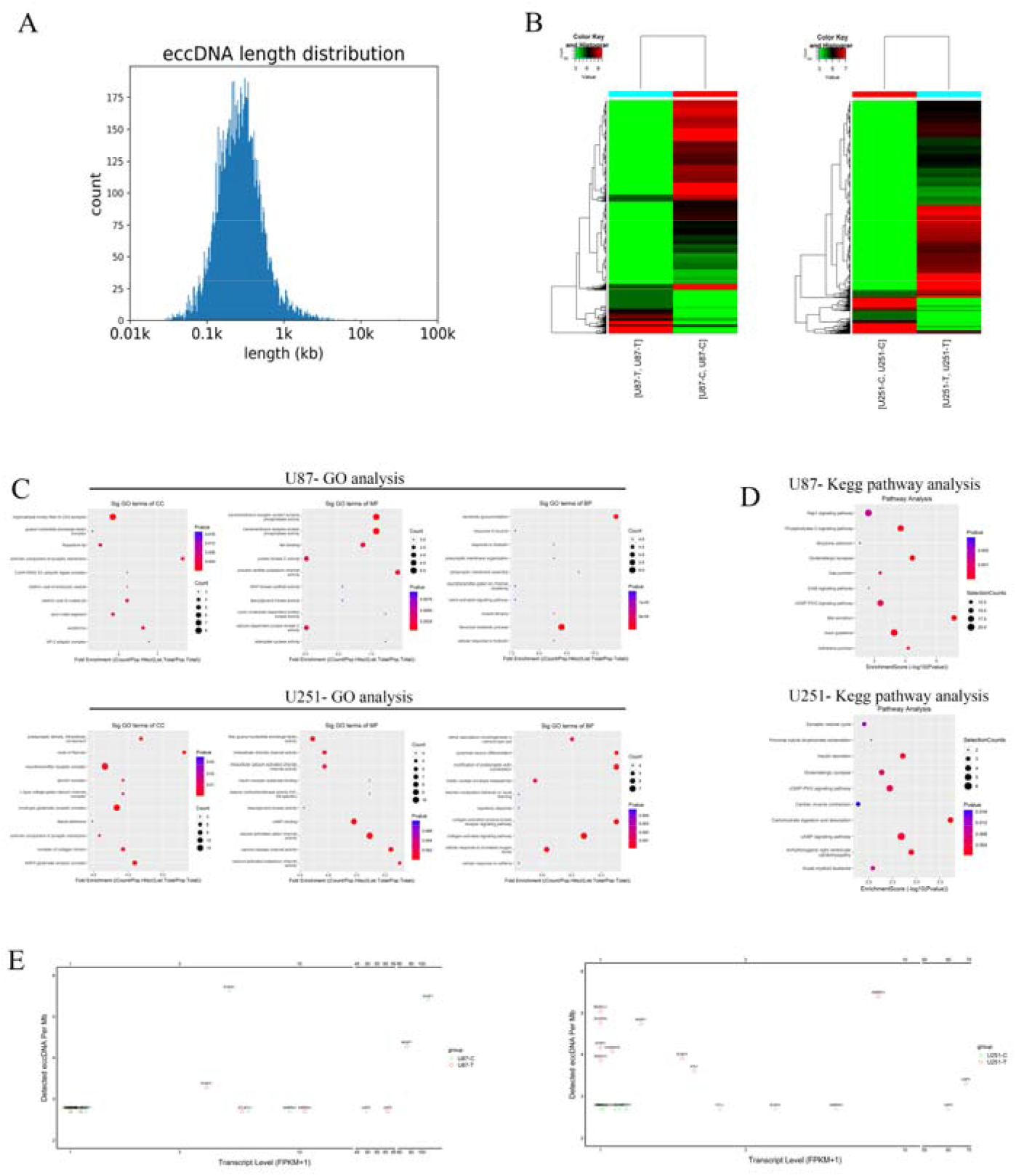
The ecDNA expression profile in U87 and U251 cells after treatment with 50μM DHA for 24h. **(A)** Extrachromosomal circle DNA length distribution of U87 and U251 cells. **(B)** Cluster analysis of ecDNA of U87 and U251 cells in both the DHA-treated and -untreated group. **(C)** GO analysis of ecDNA of U87 and U251 cells in both the DHA-treated and -untreated group. **(D)** KEGG pathway analysis of ecDNA of U87 and U251 cells in both the DHA-treated and -untreated group. **(E)** The relationship between ecDNA/MB and encoding gene/MB of U87 and U251 cells in both the DHA-treated and -untreated group.

The results of GO analysis were divided into Cell Component (CC) terms, Molecular Function (MF) terms and Biological Process (BP) terms. In the U87 cell line, after DHA treatment, the most upregulated ecDNA was enriched in synapse (CC terms), cytoskeletal protein binding (MF terms), and developmental process (BP terms). The down-regulated ecDNA was most enriched in the nodes of Ranvier (CC terms), ion binding (MF terms), and nervous system development (BP terms). In the U251 cell line, DHA treatment-upregulated ecDNA was most enriched in the extrinsic components of the synaptic membrane (CC terms), ion binding (MF terms), and the negative regulation of hippo signaling (BP terms). Downregulated ecDNA was most enriched in the cytoplasm (CC terms), vytoskeletal protein binding (MF terms), and the negative regulation of cytosolic matrix junction organization (BP terms) (Figure 2C). Further KEGG pathway analysis suggested that the upregulated ecDNA-encoding genes were most significantly enriched in the regulation of lipolysis in the adipocyte-related signaling pathway, while the downregulated genes were most significantly enriched in the axon guidance-related signaling pathway after DHA treatment in U87 cells. The genes encoding ecDNA that were upregulated by DHA treatment were most significantly enriched in the axon guidance-related signaling pathway, while the genes that were downregulated were most significantly enriched in the glutamatergic synapse-related signaling pathway in U251 cells (Figure2 D). In U87 and U251 cells, the relationship between the transcript levels and the amount of ecDNA of EMT-related genes was analyzed. We found that there was a positive correlation between ecDNA/MB and encoding gene/MB after DHA treatment, which was particularly significant in U87 cells. This suggests that ecDNA is also transcriptionally active and involved in the expression of EMT-related genes in gliomas, in addition to chromosomal linear DNA (Figure2 E)

### 3.3 Further bioinformatic analysis for potential targets based on the ecDNA expression profile and TCGA database

We acquired the RNA sequencing data and related clinical information of 701 glioma samples and 150 normal samples from the TCGA database. Through differential analysis based on RNA-sequencing of glioma with logFC filter = 1 and FDR filter = 0.05, we found 13,311 significant genes which involved 297 differential parental genes of significant ecDNAs treated with DHA (Figure3 A). Next, 200 EMT-related markers from the GESA database were put into the same differential analysis to discover the 97 significant EMT markers in glioma (Figure3 B). By means of survival analysis, 242 of the 297 ecDNA parental genes were found to have significant survival differences, as did 88 of the 97 EMT-related genes. The Venn diagram demonstrated the intersections among differential genes in glioma, significant parental genes of ecDNA, and EMT markers, which presented six targets including BASP1, CALD1,ITGA2,LAMA2,FAP,COL11A1 (Figure3 C). These six markers were put into univariate COX regression analysis, and all six of these genes were found to have an impact on the prognosis of patients. To exclude confounding factors, we conducted multivariate COX regression analysis, generating the Risk Score model with BASP1, CALD1, ITGA2 and LAMA2 (Figure3 D). The regression equation was: Risk score= −0.0087 * Expression (BASP1) + 0.0277 * Expression (CALD1) + 0.08 * Expression (ITGA2) + 0.0548 * Expression (LAMA2). The ROC curve was drawn to prove the favorable diagnostic value of the COX regression model, with an area under the ROC curve (AUC) of 0.771 (Figure3 E). We analyzed the survival status of the glioma patients from the TCGA database (Figure3 F) and divided them into a high-risk group and a low-risk group, according to the COX regression model (Figure3 G). Some of the patients from the TCGA database were assigned to the training set where the COX regression model was generated, while the other patients were assigned to the test set. Survival analysis was performed, respectively, in both sets, suggesting that the patients in the high-risk group have poor prognosis (Figure3 H-I). Heatmap analysis demonstrated that the expression of BASP1 was lower in the low-risk group than in the high-risk group, while the other three genes had a contrary tendency (Figure3 J). Survival analysis also indicated that patients’ survival time was negatively related to BASP1 level (Figure3 K). These results suggest that BASP1 might be a significant cancer suppressor gene in glioma.

**Figure 3.**
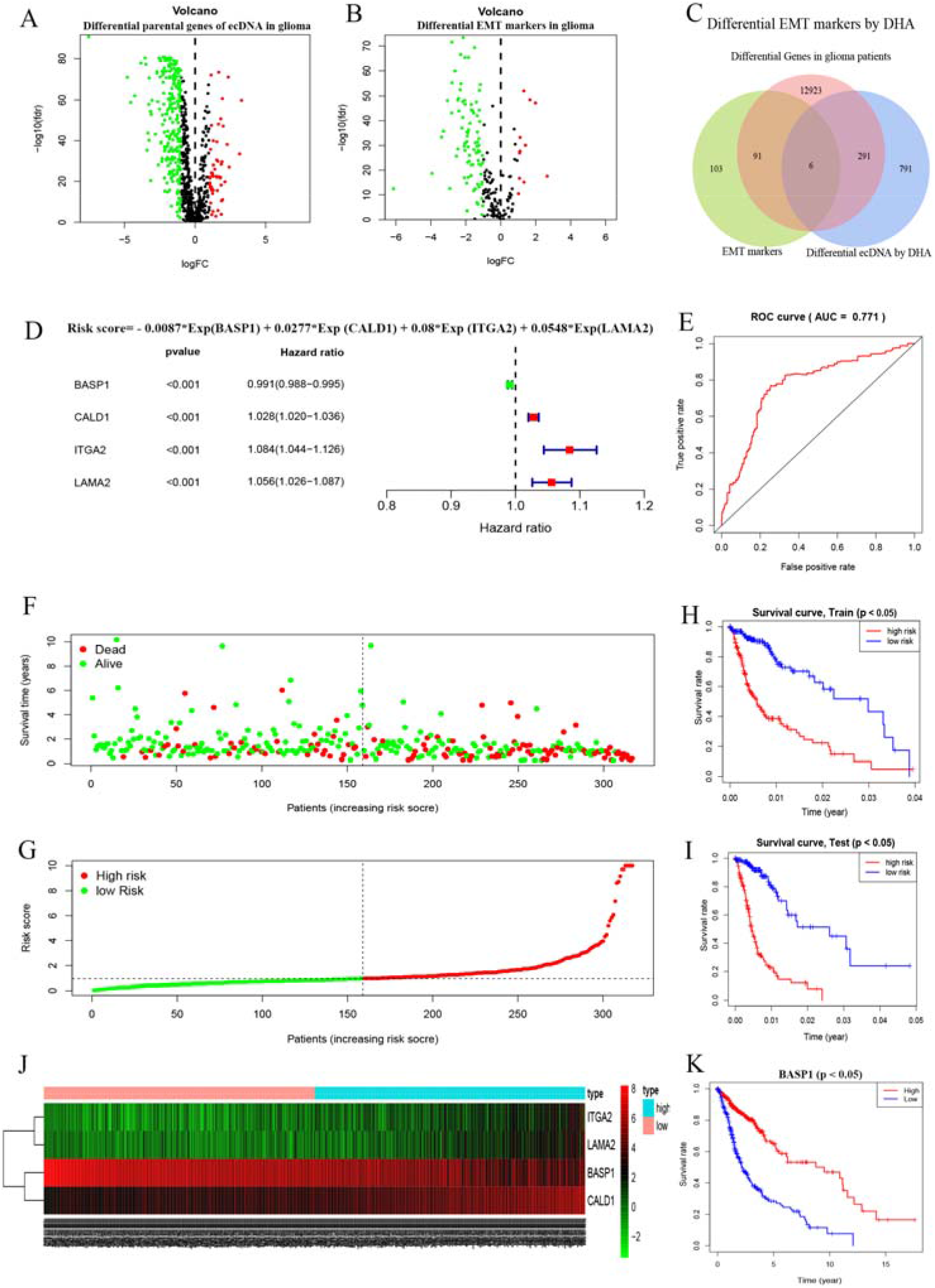
Bioinformatic analysis and target profiles. **(A)** Volcano plot for significant parental genes of ecDNA treated with DHA based on glioma samples from the TCGA database. **(B)** Volcano plot for EMT markers based on glioma samples from the TCGA database. **(C)** Venn diagram showing the intersection of significant parental genes of ecDNA and differential EMT markers. **(D)** COX regression model based on the intersecting markers. **(E)** Receiver operator characteristic curve estimating the favorable diagnostic value of the COX regression model. **(F)** The overall survival status of the patients from the TCGA database. **(G)** The patients were divided into a high-risk group and low-risk group based on the Risk Score from the COX regression model. **(H)** Survival analysis between the high-risk group and the low-risk group among patients in the training set. **(I)** Survival analysis between the high-risk group and the low-risk group among patients in the test set. **(J)** Heatmap analysis for the four targets between the high-risk group and the low-risk group. **(K)** Survival analysis for BASP1 level.

### 3.4 Confirm the level of the ecDNA from BASP1 in glioma cell lines and tissue samples

The expression of ecDNA-BASP1 was significantly up-regulated in the ecDNA expression profile of U87 and U251 cells treated with DHA. To verify the accuracy of sequencing, we used real-time PCR and Sanger sequencing to validate ecDNA-BASP1. The PCR amplification results suggest that the expression of ecDNA-BASP1 was significantly increased in U87 and U251 cells after 50μM DHA treatment for 24 h (Figure4 A). Sanger sequencing results suggested that the exact location of ecDNA-BASP1 was chr5:17125366-17127290 (Figure4 B).

**Figure 4.**
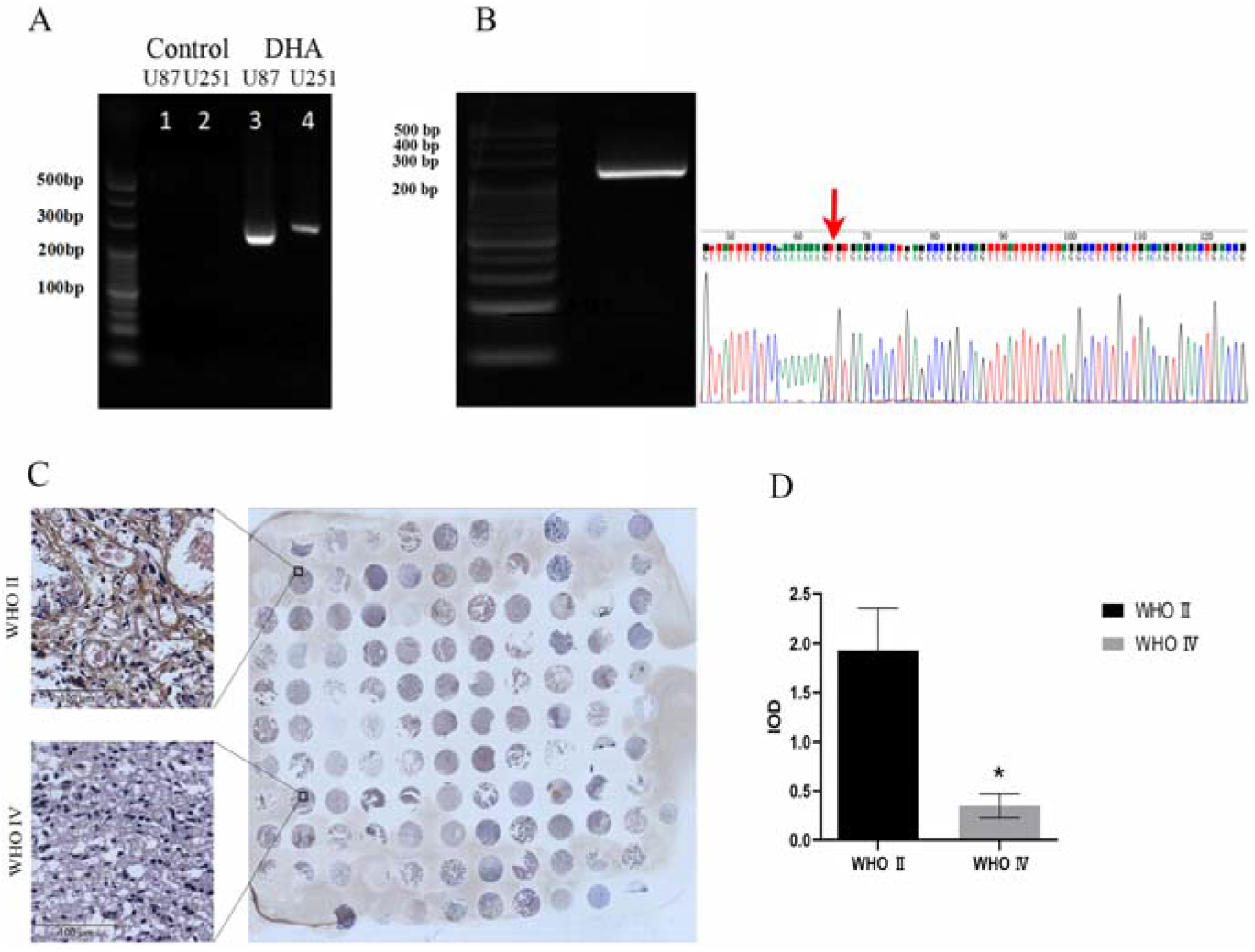
Target ecDNA gene selection in ecDNA. **(A)** Verification of ecDNA-BASP1 by PCR. **(B)** Verification of ecDNA-BASP1 by Sanger sequencing. **(C)** The relationship between ecDNA-BASP1 and glioma grade from the in situ hybridization assay. **(D)** Statistical results of the in situ hybridization assay.

To investigate the role of ecDNA-BASP1 in glioma, we applied tissue microarrays containing 60 glioma samples of different grades. In situ hybridization probes for ecDNA-BASP1 were synthesized based on the gene sequence of BASP1. The in situ hybridization results showed a significant negative correlation between ecDNA-BASP1 and glioma grade (Figure 4C-D). These results suggest that BASP1 might play a role as a tumor suppressor in the process of DHA anti-glioma EMT.

### 3.5 The inhibitory effect of BASP1 on EMT in U87and U251 cells

To further investigate the function of BASP1, we first constructed BASP1 knockdown U87 and U251 cells using the CRISP/Cas9 technique, whose small guide RNA (sgRNA) was confirmed by Sanger sequencing (Figure5 A). The CCK-8 assay revealed that the proliferation ability of U87 and U251 cells was significantly increased after knockdown of BASP1 (Figure5 B-C), suggesting that BASP1 has an inhibitory effect on the proliferation of U87 and U251 cells. The cell scratch assay revealed that the migration ability of U87 and U251 cells was enhanced after knockdown of BASP1 (Figure5 D-F), suggesting that BASP1 could inhibit the movement of tumor cells in U87 and U251 cells by means of some mechanism. The transwell assay further suggested that the invasive ability of U87 and U251 cells was also improved after knockdown of BASP1 (Figure5 G-L). The expression of both vimentin and N-cadherin, the proteins that promote EMT, was upregulated in the knockdown group (Figure5 M-O), suggesting that in U87 and U251 cells, BASP1 may exert its inhibitory effect on migration invasion by inhibiting EMT.

**Figure 5.**
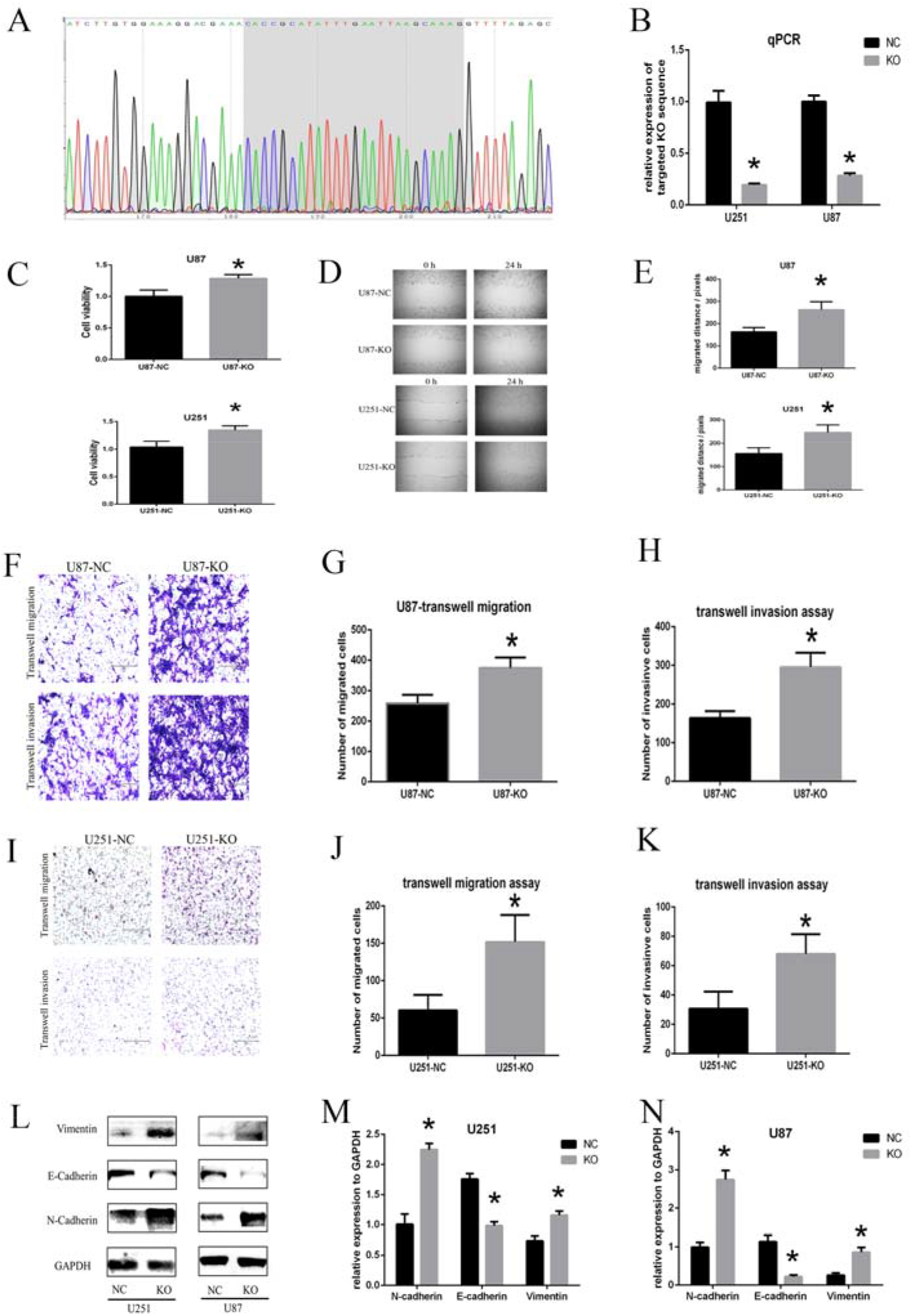
ecDNA-BASP1 inhibited migration and invasion through EMT in U87 and U251 cells. **(A)** Sanger sequencing result of the ecDNA-BASP1 KO target. **(B)** qPCR results of the targeted KO sequence. **(C)** Cell viability in U87 and U251 cells after ecDNA-BASP1 knockdown. **(D)** Wound scratch assay in U87 cells and U251 cells after ecDNA-BASP1 knockdown. **(E)** Statistic result of wound scratch assay in U87 and U251 cells after ecDNA-BASP1 knockdown. **(F)** Transwell assay showed the change of migration and invasion in U87 cells after ecDNA-BASP1 knockdown. **(G)** Statistic result of transwell migration assay in U87 cells. **(H)** Statistic result of transwell invasion assay in U87 cells. **(I)** Transwell assay showed the change of migration and invasion in U251 cells after ecDNA-BASP1 knockdown. **(J)** Statistic result of transwell migration assay in U251 cells. **(K)** Statistic result of transwell invasion assay in U251 cells. **(L)** The expression of vimentin, E-cadherin and N-cadherin in U87 and U251 cells after ecDNA-BASP1 knockdown. **(M)** Statistic result of Western blot in U251 cells. **(N)** Statistic result of Western blot in U87 cells. *P<0.05.

### 3.6 Verifying the function of ecDNA-BASP1 in vivo

We investigated the function of ecDNA-BASP1 in vivo using an intracranial in situ transplantation model in nude mice. It was found that the survival of nude mice after DHA administration was significantly longer than that of the Control group (Figure6 AB). DHA inhibited the percentage of Ki76 positive cells and increased the expression of Caspase 3 in the U87-and U251-induced orthotopic implantation model (Figure6 C-F). Analysis of ecDNA-BASP1 in intracranial tumor tissues of nude mice using the in situ hybridization technique revealed that ecDNA-BASP1 in the DHA group was significantly higher than that in the Control group, suggesting that DHA may inhibit glioma development by increasing ecDNA-BASP1 levels in U87 and U251 (Figure 6GH).

**Figure 6.**
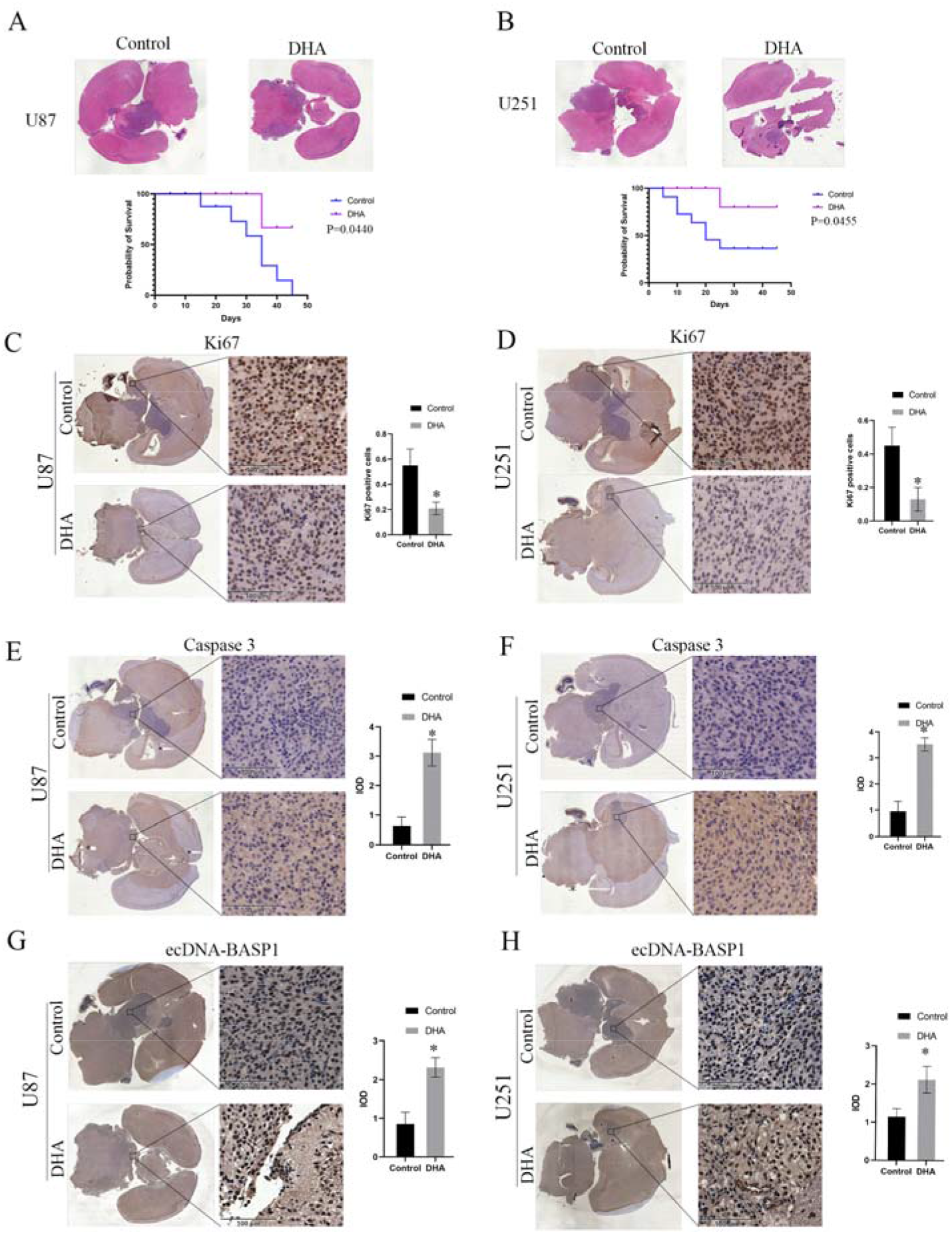
Orthotopic implantation model in nude mice. **(A)** DHA inhibited glioma growth and improved survival time in the U87-induced orthotopic implantation model, p<0.05, n=8. **(B)** DHA inhibited glioma growth and improved survival time in the U251-induced orthotopic implantation model, p<0.05, n=8. **(C)** IHC analysis of Ki67 in vivo in the U87-induced model, p<0.05, n=8. **(D)** IHC analysis of Ki67 in vivo in the U251-induced model, p<0.05, n=8. **(E)** IHC analysis of Caspase 3 in vivo in the U87-induced model, p<0.05, n=8. **(F)** IHC analysis of Caspase 3 in vivo in the U251-induced model, p<0.05, n=8. **(G)** The expression of ecDNA-BASP1 in vivo in the U87-induced model, p<0.05, n=8. **(H)** The expression of ecDNA-BASP1 in vivo in the U251-induced model, p<0.05, n=8.

## 4. Discussion

In this study, the effects of DHA on the proliferation, apoptosis, oxidative stress and migratory invasion of U87 and U251 cells are demonstrated. DHA can inhibit proliferative activity, increase ROS levels and promote apoptosis in U87 and U251 cells. The migratory invasion activity can also be effectively inhibited. EcDNA sequencing was applied to explore the changes to ecDNA expression in U87 and U251 cells after the treatment with DHA. By analyzing the ecDNA expression profiles of U87 and U251 cells, we found that most ecDNA expression was positively correlated with transcriptional activity, indicating that most ecDNA was involved in the transcription of genes. Combined with the survival analysis, we finally selected ecDNA-BASP1 as the target involved in DHA inhibition of EMT in U87andU251 cells. By means of tissue microarray in situ hybridization experiments, ecDNA-BASP1 was found to be present at all levels of GBM tissues and positively correlated with patient prognosis. It is suggested that BASP1 may play a role as an oncogene inhibitor in GBM. After knockdown of BASP1 using CRISP/Cas9, we found that the proliferation of glioma cells was significantly enhanced, and migration and invasion were upregulated, consistent with a suppressor gene feature. We subsequently found that DHA was able to inhibit the growth of intracranial transplantation glioma in nude mice. In situ hybridization suggested that the ecDNA-BASP1 content in nude mouse transplant tumors was significantly increased after DHA treatment, which might exert a better suppressive effect on glioma.

Several studies have found that DHA can effectively inhibit the proliferation of glioma cells and promote oxidative stress and apoptosis production [23,24]. In this study, the above findings were first verified in U87 and U251 cells. The inhibitory effect of DHA on the migration and invasion of GBM and the related molecular mechanisms have also been reported in recent years, as Hou, Kuiyuan et al. found that DHA could inhibit the migration and invasion of glioma through NHE1 and other mechanisms [25]. In this study, the inhibitory effect of 50μM DHA on the migration and invasion of U87 and U251 cells was subsequently verified by means of transwell assay and ibidi Culture-Insert 2 Well. There is increasing evidence that ecDNA plays an important role in tumor biology. Considering that unlike chromosomal linear DNA in the nucleus, ecDNA is present in the cytoplasm and may be more susceptible to ROS, we examined the changes in ecDNA in U87 and U251 cells due to the effect of DHA. The results show that ecDNA was present in large amounts in U87 and U251 cells in both DHA-treated and DHA-untreated group and the fragments were distributed between 0.01 and 10 kb. GBM was found to be the tumor with the highest ecDNA content, and most of the current basic studies on ecDNA have used cell lines derived from GBM [26]. Analysis of the changes in ecDNA expression profiles of U87 and U251 cells in both the DHA-treated and DHA-untreated group revealed that a large number of ecDNA can carry coding genes including oncogenes and suppressor genes. In previous ecDNA studies, researchers have focused more on oncogenes carried by ecDNA or on phenotypes that favor tumor cell survival such as tumor drug resistance, and suggested that tumor cells could evolve faster and adapt to the microenvironment through ecDNA mechanisms [27]. Notably, the expression of some antioncogenes in the ecDNA expression profile was also elevated after the effect of DHA in our study. By reviewing the literature, we speculated that this may be related to the mechanism of ecDNA production. In tumor cells, ecDNA production is associated with chromosome fragmentation and is regulated by poly ADP-ribose polymerases (PARP) and DNA-dependent protein kinase catalytic subunits (DNA-PKcs) [28]. As shown in Figure 7, we hypothesized that the oxidative stress triggered by DHA treatment could cause DSBs extensively and indiscriminately in a short period of time, exacerbating chromosome fragmentation. This prevents tumor cells from increasing only the oncogenic ecDNA mediated by potential regulatory mechanisms, and the ecDNA carrying suppressor genes will also increase and eventually participate in the transcriptional translation of a suppressor gene to exert subsequent effects. We selected ten genes related to EMT and migratory invasion by bioinformatics analysis, and found that the amount of ecDNA was positively correlated with the transcriptional activity of suppressor genes, suggesting that the inhibitory effect of DHA on the migratory invasion of U87 and U251 cells is partly achieved by changing ecDNA.

**Figure 7.**
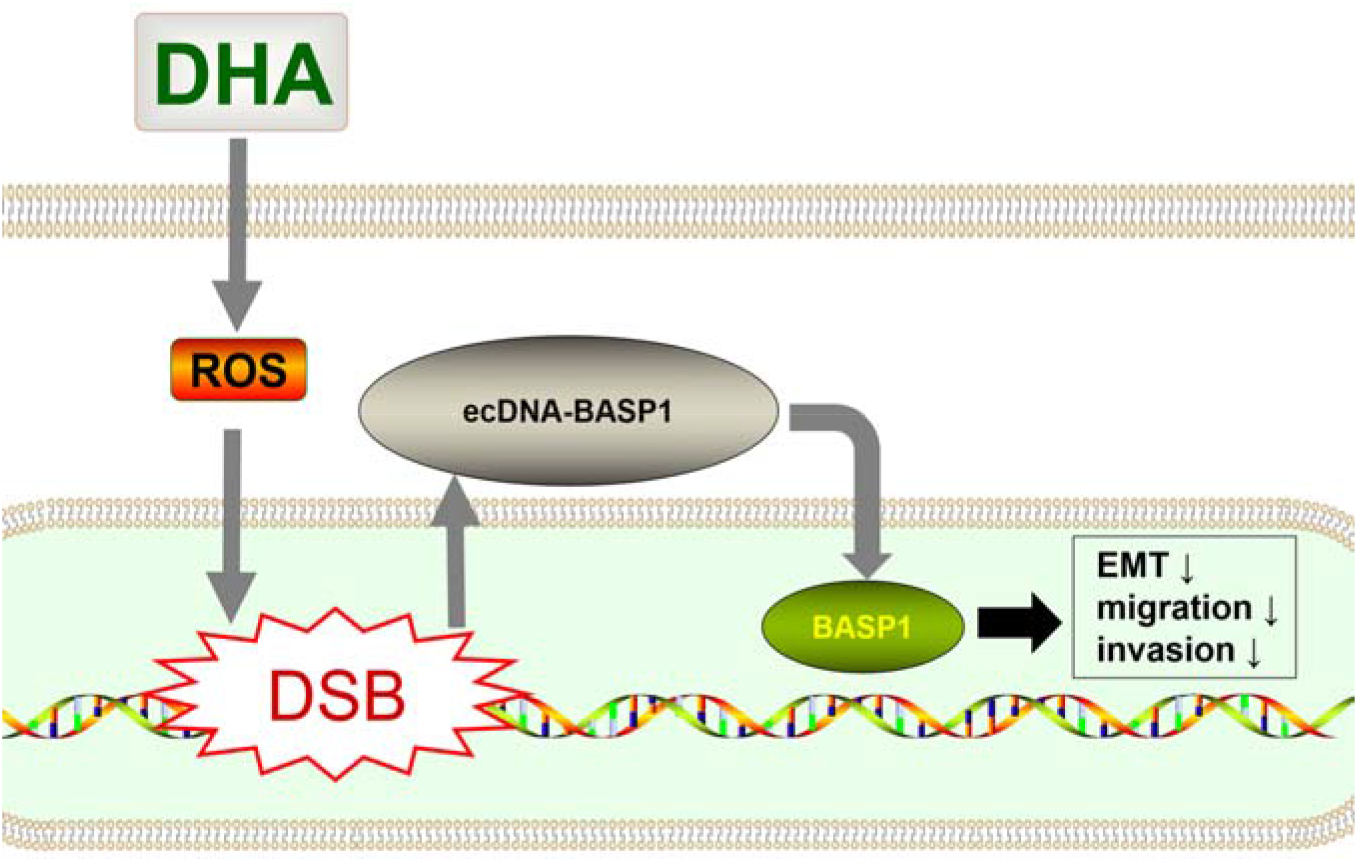
Schematic illustration of DHA inhibiting EMT through ecDNA-BASP1.

Through bioinformatic analysis, we finally focused on ecDNA carrying brain abundant membrane attached signal protein 1 (BASP1), which is a member of the neural signaling protein family along with myristoylated alanine rich protein kinase C substrate (MARCKS) and growth-associated protein 43 (GAP43). MARCKS and GAP43 are members of the neural signaling protein family, and they share the same structural domain with the intracellular membrane lipid component phosphatidylinositol (4,5) bisphosphate (PIP2), which binds to the inner side of the cell membrane. It binds to calmodulin (CaM), actin and is regulated by protein kinase C to regulate neural cell growth [29]. In addition to neural tissues, BASP1, MARCKS and GAP43 are also expressed. The current study found that in tumors, MARCKS and GAP43 have oncogenic effects, while BASP1 exerts anti-oncogenic functions [30–32]. Unlike in neuronal cells, in tumor cells, BASP1 protein is translocated to the nucleus to function as a transcription factor for regulatory purposes. The present study confirmed that the most important anti-cancer mechanism of BASP1 is its binding to Williams’ tumor 1 (WT1) as a complex to act as a repressor of oncogene transcription [33]. In a follow-up study using an in situ hybridization assay on tissue microarrays of 60 GBM cases, we found that ecDNA-BASP1 content was negatively correlated with WHO grading in patients with different grades of GBM. It is suggested that BASP1 protein may function as a tumor suppressor gene in GBM, a finding which is consistent with previous studies [32]. To further investigate the function of BASP1, we knocked down the BASP1 gene in U87 and U251 cell lines by CRISP/Cas9. The results showed that the proliferation ability and migration invasion ability of U87 and U251 cells were improved after knockdown of BASP1. Then, the orthotopic implantation model in nude mice verified that ecDNA-BASP1 expression increased after DHA treatment. These results further confirmed that DHA could inhibit the migration invasion of U87 and U251 cells by upregulating ecDNA-BASP1.

In this study, we firstly confirmed the role of ecDNA in DHA-inhibiting glioma. EcDNA in U87 and U251 cells was investigated for the first time and preliminary bioinformatic analysis of expression profiles during DHA treatment was performed. For the first time, DHA was found to be able to finally inhibit the migration invasion of U87 and U251 cells through the upregulation of ecDNA-BASP1. The above findings have never been reported.

However, there are still some shortcomings in this study that need to be further investigated in the future. First, this study only found that DHA can upregulate a portion of ecDNA, and the specific molecular mechanism is not clear, which may be related to the more fundamental biological mechanism of ecDNA production and degradation. Secondly, the specific mechanism by which ecDNA-BASP1 inhibits the migratory invasion of U87 and U251 needs to be further explored. Additionally, some studies have shown that BASP1 can directly inhibit the transcriptional activity of WT1 after binding to WT1[34]. Additionally, some studies have found that Yes-associated protein (YAP) can bind to WT1 and inhibit the expression of E-cadherin expression, which triggers the EMT process and thus promotes cell motility [35]. Therefore, we can speculate that BASP1 may inhibit the YAP-WT1-mediated down-regulation of E-cadherin expression by competing with WT1 to suppress the EMT of glioma cells and thus inhibit migration and invasion. In addition, ecDNA sequencing in this study was limited to the analysis of coding genes only, and the analysis of non-coding genes was not included. Therefore, the miRNA, lnRNA and circRNA genes carried by ecDNA should be studied systematically in future studies.

## 5. Conclusions

DHA was able to inhibit the proliferation, apoptosis, migration, and invasion activities of U87 and U251 cells. The ecDNA expression profile in U87 and U251 cells was dramatically altered by DHA. Among them, the upregulation of antioncogene ecDNA-BASP1 expression played an important role in the inhibition of glioma growth and infiltration by DHA. ecDNA-BASP1 may inhibit the migration and invasion of glioma by suppressing EMT.

## Supplementary Materials

All the necessary supplementary materials have been uploaded.

## Author Contributions

Conceptualization, Que Zhongyou and Lei Bingxi; methodology, Lei Bingxi; software, Lei Bingxi.; validation, Zhou Zhiwei; formal analysis, Lei Bingxi; data curation, Que Zhongyou and Lei Bingxi; writing—original draft preparation, Que Zhongyou; writing—review and editing, Lei Bingxi and Zhou Zhiwei; visualization, Lei Bingxi; supervision, Liu Sheng, Zheng Wenhua; project administration, Que Zhongyou; funding acquisition, Que Zhongyou and Liu Sheng. All authors have read and agreed to the published version of the manuscript.

## Funding

Please add: This research was funded by Shenzhen Science and Technology Innovation Commission(JCYJ20180307155043326).

## Institutional Review Board Statement

The animal study protocol was approved by the Animal Experiment Ethics Committee of Sun Yat-Sen University (LACUC number: L102022016000D). All patients provided informed consent for clinical research.

## Data Availability Statement

The data presented in this study are available on request from the corresponding author.

## Acknowledgments

We thank Zhang longping, Li weixin and Yuan Bo for their great help in our research.

## Conflicts of Interest

The authors declare no conflicts of interest.

## Notes

### Competing Interest Statement

The authors have declared no competing interest.

### Summary of Updates

The order of authors has been corrected.

